# Community-level diversity of viral counter-defensomes

**DOI:** 10.64898/2026.08.10.744029

**Authors:** Angelina Beavogui, Lucas da Silva e Silva, Izabel de G. M. Gomes, Luke Doladille, Nicolas Wiart, Patrick Wincker, Pedro H. Oliveira

**Author notes:** These authors contributed equally to this work (Angelina Beavogui, Lucas da Silva e Silva).

## Abstract

Bacteria and phages have co-evolved tit-for-tat strategies to combat one another for survival. To overcome phage infection, bacteria have developed various defense systems (the defensome), and in response, phages have evolved a counter-defensome to inhibit or evade such defensive strategies. While significant progress has been achieved on mechanistic, structural and functional aspects of the defensome, the study of the phage counter-defensome is relatively new, with very limited information available on the diversity and distribution of these systems across complex environmental phageomes. Here we present a large-scale analysis of the counter-defensome of 50,595 DNA phage population genomes reconstructed from soil, marine, and human gut environments. We observed substantial variation in the frequency and composition of the counter-defensome across phage families, which correlated with host range, habitat, and geographic context. A substantial fraction of the counter-defensome was clustered in islands, with gene expression restricted to a limited subset of families. Moreover, both phage counter-defense and bacterial-type defense families were detected across genomes of nucleocytoplasmic large DNA viruses, suggesting cross-clade horizontal gene transfer. Our results reveal the diverse counter-defense strategies across environmentally distinct viral communities and provide a foundation for uncovering novel mechanisms that shape conflicts and alliances in host–virus immunity networks.

## Introduction

Viruses are the most abundant, pervasive, and genetically diverse biological entities on Earth. Among them, bacteriophages (phages) are profoundly influential in shaping microbial ecosystem dynamics, biogeochemical nutrient cycling, and gene flow through horizontal gene transfer (HGT). With an estimated global population size of 10^31^ particles ^1^, phage abundance relative to their hosts, often quantified as a virus-to-prokaryote ratio (VPR), spans more than six orders of magnitude across environments as disparate as the human gut, the global ocean, and soils ^2,3^. Most known prokaryotic viruses are tailed double-stranded (ds) DNA viruses pertaining to the class *Caudoviricetes* (realm *Duplodnaviria*), with additional modest contributions from *Varidnaviria* and archaeal viruses of *Adnaviria* ^4,5^. They typically possess genomes ranging from tens to hundred of kilobases, with characteristically mosaic architectures, diverse replication strategies, and variable host ranges. Beyond prokaryotes, certain dsDNA viruses can also infect unicellular eukaryotes. An interesting example is provided by virophages (class *Maveriviricetes*), which parasitize giant viruses of the order *Imitervirales* (phylum *Nucleocytoviricota*) also known as nucleocytoplasmic large DNA viruses (NCLDVs) ^6^. Virophages depend on the viral factory provided by the co-infecting NCLDV for their own replication, eventually impairing the latter’s fitness and benefiting the host cell ^7^. Comparatively less studied than phages, virophages and NCLDVs appear to comprise only a small fraction of the virosphere (often less than 1% of the viral reads in environmental metagenomes) ^8–10^, although NCLDVs can reach 10^3^–10^5^ genomes per milliliter in aquatic ecosystems ^11^. At evolutionary time scales, dsDNA viruses have the capacity to mediate gene transfer across deep evolutionary lineages, as previously reported for NCLDVs and eukaryotes ^12^, and potentially between cellular domains through indirect or mobile genetic element (MGE)-mediated mechanisms ^6,12,13^.

This pervasive viral pressure has driven bacteria to evolve a sophisticated repertoire of defense systems, collectively termed the defensome ^14,15^. Classical examples include restriction–modification (R–M) systems ^16^ and CRISPR/Cas adaptive immunity ^17^, which detect and degrade invading nucleic acids. Beyond these well-characterized mechanisms, the past decade has revealed a vast arsenal of previously unknown anti-MGE defense systems, many of which have begun to be mechanistically characterized ^18,19^. Phages, in turn, have responded by evolving a growing array of counter-defense strategies capable of targeting bacterial defensomes at multiple levels, with more than 150 systems and 27 families described to date ^20^. These include direct inhibition of defense proteins (*e.g.*, anti-CRISPR (Acr) ^21,22^, anti-RecBCD ^23^, anti-restriction enzymes ^24,25^), interference with toxin–antitoxin (anti-T–A) systems ^26,27^), sequestration of immune signaling molecules (*e.g.*, anti-CBASS ^28,29^, anti-Thoeris ^30^), and chemical modification of host defense components ^31,32^. Apart from counter-defenses, phage genomes, particularly in the lysogenic state, display pronounced genomic mosaicism and often carry anti-MGE genes implicated in inter-viral competition, capable of increasing host resistance against infection from competing phages ^33,34^. Such layer of accessory ‘moron’ genes comprising defense and counter-defense is often acquired and lost via homologous recombination and lateral transduction ^35^. Arms races are also present among eukaryotic viruses. For example, giant viruses have evolved defense systems to counter virophage parasitism, mirroring bacterial anti-MGE strategies. The best-characterized example is MIMIVIRE, a CRISPR-like adaptive immune system identified in mimiviruses, which incorporates virophage-derived sequences to mediate sequence-specific resistance ^36^. Emerging evidence suggests that giant viruses encode a broader repertoire of virophage defense mechanisms, including DNA modification and restriction nucleases ^37^, underscoring evolutionary parallels between prokaryotic and viral immunity.

Recent advances in high-throughput sequencing and computational metagenomics have begun to illuminate largely hidden dimensions of viral biology. Enhanced metavirome extraction, combined with culture-independent, genome-resolved approaches, now enables systematic exploration of the vast viral ‘dark matter’ across diverse ecosystems ^38–42^, providing opportunities to study its abundance and diversity across environments. These advances reveal pronounced temporal and spatial variation in VPRs, suggesting that fluctuating viral predation exerts a major influence on the composition and evolution of immune repertoires across biomes. Here, we systematically characterized the counter-defensome of dsDNA phage communities across soil, marine, and human gut ecosystems. We quantified its abundance, diversity, and genomic organization, examined patterns of co-localization within counter-defense islands, and assessed associations with host taxa and biogeography. We further mapped the landscape of bacterial anti-MGE defense genes carried by phages, and the overall expression repertoire of phage immune modules. Finally, we extended our analysis to virophages and NCLDVs, revealing a continuum of conserved strategies across viral lineages. Together, our findings contribute to a better understanding of the complexity of viral immune networks and highlight their central role in shaping microbial community structure and evolutionary dynamics across the biosphere.

## Results

### Abundance and distribution of counter-defense elements in phageomes

We first examined the distribution of phage-encoded counter-defense elements across a comprehensive collection of 50,595 high-quality viral operational taxonomic units (vOTUs, *≥*90% completeness and *≤*5% contamination or redundancy, see Methods) recovered from soil, marine, and human gut environments (Fig. 1a, Supplementary Data 1–2, Supplementary Fig. 1a). In parallel, we extended our analysis to identify and quantify phage-encoded anti-MGE systems of bacterial origin, which are typically co-opted for inter-MGE conflicts ^43^. Throughout this study, we considered a counter-defense or anti-MGE system to be complete when it retains an experimentally validated genetic architecture sufficient to confer immune activity. By contrast, individual counter-defense and anti-MGE genes are interpreted as remnants of these systems, arising either through HGT or through the progressive decay of once-intact loci. The composition of our vOTU collection is summarized in the Sankey diagrams in Fig. 1b, which connect phage morphology, family- and subfamily-level taxonomy, and sample provenance across the three environments. As expected given their global prevalence, head-tailed phages of the class *Caudoviricetes* predominate in all datasets, with the *Siphoviridae*, *Podoviridae*, and *Autographiviridae* families accounting for most recovered vOTUs. Across specific environments, soil vOTUs are dominated by *Siphoviridae* and uncategorized lineages, marine vOTUs are more evenly distributed among multiple families across ocean regions, while human-gut viromes exhibit the highest overall family richness, with *Siphoviridae* as the single most represented group across geographically diverse cohorts.

**Fig. 1.**
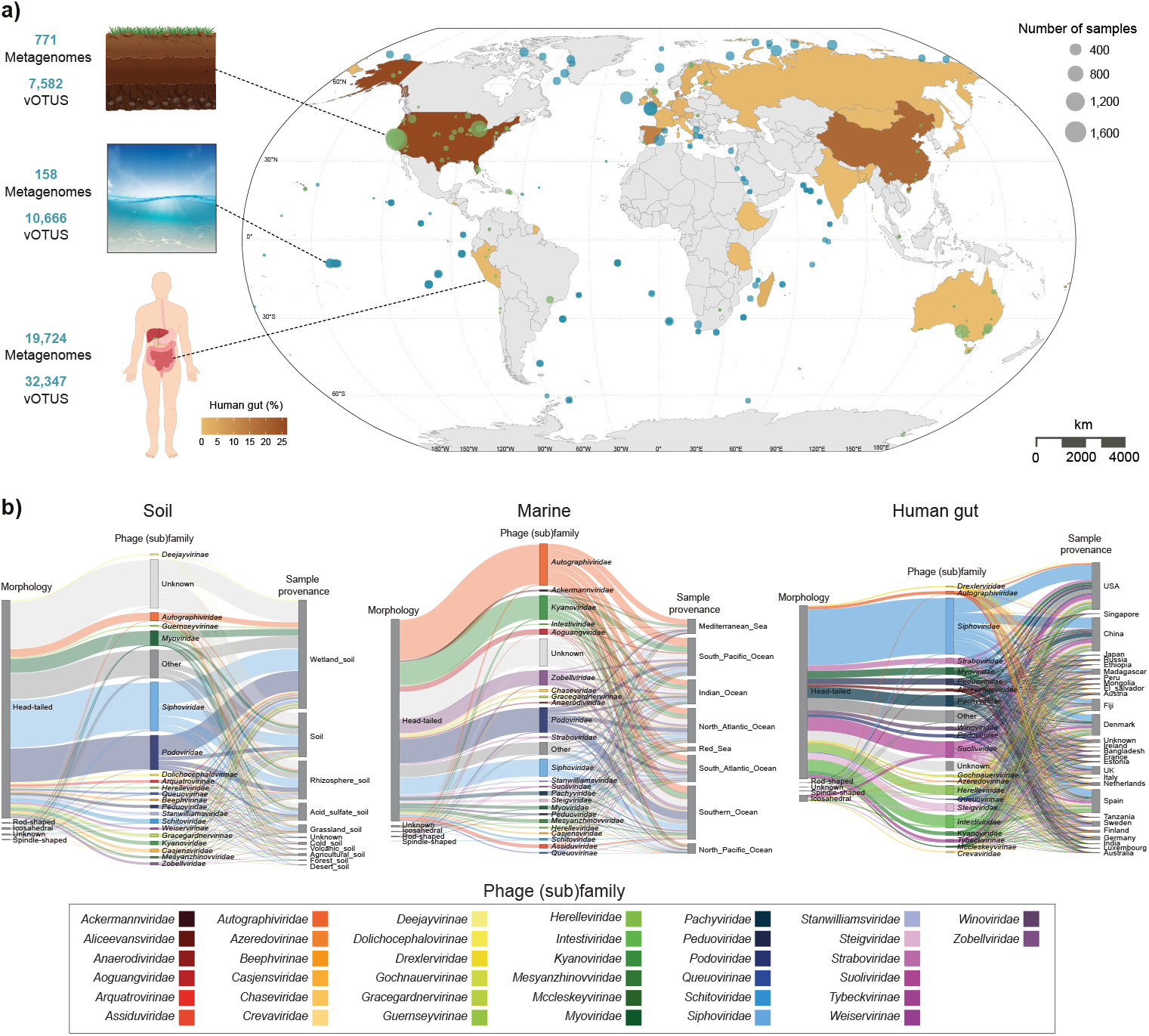
Dataset characterization. **a**) Our analysis comprised 50,595 high-quality viral operational taxonomic units (vOTUs) recovered from 20,653 metagenomes spanning three major environments: soil, marine, and the human gut. The geographic distribution of marine and soil sampling sites is shown by latitude and longitude coordinates on the world map, alongside a heatmap indicating the proportion of human gut samples contributed by each country. **b**) Sankey diagrams illustrating the relationships among phage morphology, family and subfamily-level taxonomy, and sample provenance. Image credits (copyright-free) for panel **a**): soil (brgfx/Freepik), sea (vectorpocket/Freepik), and body/intestines (brgfx/Freepik).

In our vOTU dataset, we identified a total of 15,080 counter-defense systems and 17,756 counter-defense genes spanning 18 counter-defense families (Fig. 2a, Supplementary Data 3, Supplementary Fig. 1b). The relative distribution of counter-defense systems varied markedly across environments, with phages from the human gut exhibiting both higher diversity and greater system density per genome compared to soil and marine counterparts. Acr, anti-R–M, and anti-T–A systems emerged among the most abundant counter-defense strategies across all environments and were consistently detected in both virulent and temperate phages (Fig. 2a). This pattern mirrors the widespread prevalence of corresponding anti-MGE defensome families in bacterial genomes ^15^, suggesting a tight co-evolutionary coupling between bacterial immunity and phage countermeasures. Also consistent with trends previously reported for bacterial anti-MGE defensomes ^15^, we observed a negative correlation between counter-defensome density and medium-to-large sized phage genomes, particularly in human gut biomes (Supplementary Fig. 1c). This relationship is compatible with the notion that smaller phage genomes (typically associated with temperate lifestyles) engage more frequently in horizontal genetic exchange ^44^, and favor the acquisition and maintenance of mechanisms to evade host defenses. The number of counter-defense systems per vOTU followed a geometric-like distribution with most phages (75.1%) showing a limited arsenal of counter defense systems, and a global mean value of 0.3 (Fig. 2b). The latter is approximately ten-fold lower than the average number of defense systems reported per bacterial genome ^15^ across similar environments, likely reflecting both biological asymmetries in host–virus interactions and the current bias in our knowledge of defensomes relative to counter-defensomes. Still, a subset of human gut-associated *Myoviridae* phages displayed markedly expanded repertoires with up to 11 systems (Fig. 2b). Such outliers consist essentially in anti-R–M and likely reflect adaptation to hosts with particularly R–M-rich genomes, including species such as *Helicobacter pylori*. Stratification by phage family and subfamily revealed finer-scale patterns of counter-defensome enrichment. For example, members of *Schitoviridae* and *Podoviridae* appear to act as preferential reservoirs of specialized inhibitors, including anti-Thoeris and anti-Dnd systems, while others such as *Intestiviridae* are relatively depleted (Fig. 2c, Supplementary Data 4). Across environments, *Siphoviridae*- and *Podoviridae*-like phages encoded the broadest repertoire of counter-defense systems, including Acr, anti-R–M, anti-T–A and anti-SOS response modules (Figs. 2c,d, Supplementary Fig. 1d). *Proteobacteria* represented the predominant host phylum for soil and marine vOTUs, supporting a wide diversity of counter-defense strategies, whereas human-gut vOTUs were primarily associated with *Firmicutes* and *Bacteroidota* hosts and enriched in Acr, anti-SOS, and anti-Thoeris systems. Soil and marine environments showed a higher prevalence of counter-defense systems carried by virulent phages, whereas the human gut biome showed a higher prevalence of these systems carried temperate phages (Supplementary Fig. 1e). This pattern likely reflects: *i*) the lysogenic lifestyle of temperate phages in the human gut; and *ii*) the comparatively short and less frequent phage–bacteria contact in soil and marine environments, which favors virulent phages that carry counter-defense elements enabling them to overcome host immunity within a single infection attempt.

**Fig. 2.**
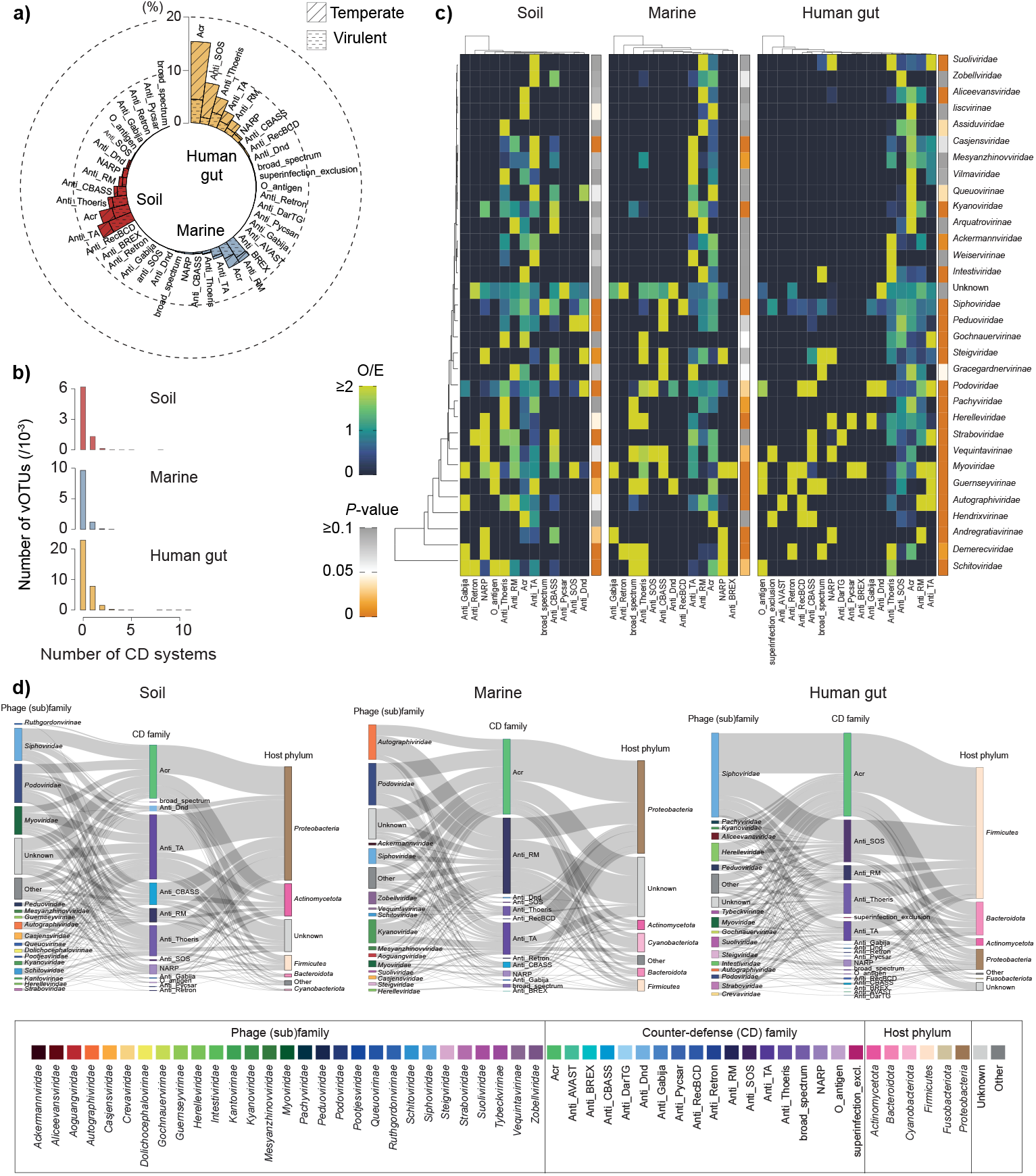
Landscape of counter-defense (CD) systems across viral populations. **a**) Proportion of soil, marine, and human gut vOTUs encoding each family of CD systems. **b**) Frequency distribution of CD system counts (per vOTU) across environments. **c**) Heatmap showing observed-to-expected (O/E) ratios for distinct CD families. Only families detected across all three environments (soil, marine, and human gut) are displayed. Expected values were calculated by multiplying the total number of systems within each counter-defense family by the overall proportion of all counter-defense-associated systems assigned to each viral (sub)family across the full dataset. Statistical significance was assessed using a two-sided Chi-square test. **d**) Sankey diagrams tracing the connections among phage family or subfamily, CD family, and host phylum across environments.

While not directly implicated in neutralizing host defense systems, we also examined the presence of anti-MGE defenses in phages, which are thought to mediate inter-MGE conflicts and thereby favor their retention within host cells. We found 2,874 anti-MGE systems and 60,145 anti-MGE genes pertaining to a total of 191 families (Supplementary Data 3). These genes proved to be unevenly distributed across viral genomes and were globally dominated by R–M, Cas and All_UG families (Supplementary Fig. 2a). Particularly for human gut biomes, smaller genomes tended to be denser (per kb) in anti-MGE defense genes (Supplementary Fig. 2b), and most phages (*∼*69%) harbored at most one anti-MGE defense gene (Supplementary Fig. 2c). At the phage-family level, the dominant families in each ecosystem carry the bulk of the anti-MGE load, with prominence in *Kyanoviridae*, *Autographoviridae* and *Siphoviridae* (Supplementary Fig. 2d). Similarly to counter-defense, soil and marine environments showed a higher frequency of anti-MGE genes in virulent phages, whereas the human gut biome showed a slightly higher prevalence in temperate phages (Supplementary Fig. 2e). No significant differences were observed in the density of anti-MGE genes after splitting the dataset in regular and jumbo phages (Supplementary Fig. 2f), suggesting a saturation effect with genome size similar to that observed in bacteria ^15^.

Overall, these results show that counter-defense and anti-MGE systems are widespread but unevenly distributed across phageomes, with clear stratification by environment, genome size and phage taxonomy. Their enrichment in specific habitats, particularly the human gut, smaller-sized genomes, and their concentration in distinct phage lineages indicate that the acquisition and maintenance of these loci are tightly linked to local host ecology and lifestyle.

### Co-localization of phage-encoded immune modules

Given the widespread distribution of counter-defense and anti-MGE systems/genes across phage genomes, we next quantified the extent of their co-occurrence, colocalization, and assessed whether they cluster into genomic islands (here termed counter-defense islands), in analogy to bacterial defense islands (Methods). Restricting the analysis to the five most abundant counter-defense and anti-MGE families, we found that genes pertaining to Dpd, Anti-Thoeris, and Anti-SOS more frequently co-occurred with other counter-defense gene families within the same phage genome (Fig. 3a). Among all pairwise combinations, Dpd/All_UG, Acr/Anti-SOS, Dpd/Anti-Thoeris, and Anti-SOS/Anti-Thoeris exhibited the highest observed-expected ratios, whereas Anti-TA/Acr and Anti-SOS/All_UG showed the lowest ones. Consistent with these findings, several immune gene families were found to co-localize (Fig. 3b), with counter-defense and anti-MGE modules frequently interspersed within the same genomic loci. In one representative example, a phage from an unknown family recovered from the human gut (uvig_530261) harbored a Type II R–M system flanked by a P1 *dar* Anti-R–M system of Type I, a *cas12f* (type V-F) effector gene, and the Anti-Thoeris *tad2* gene (Fig. 3b). This organization expands the repertoire of genomic regions that combine immune evasion and immune interference functions, extending previous observations of multi-layered counter-defense architectures ^20^. At a global scale, and irrespective of environmental context, we observed preferential co-localization between R–M / SspBCDE and to a lesser extent MADS / Acr (Fig. 3c, Supplementary Data 5). While the former suggests interplay between methylation and phosphorothioation, the latter parallels the MADS–CRISPR/Cas association observed in bacteria ^45^. In contrast, other combinations exhibited stronger environment-dependent patterns. For example, Cas / anti-Thoeris, anti-T–A / anti-Thoeris, anti-T–A / anti-CBASS co-localization was more prevalent in soil and/or human gut viral samples, whereas Gabija / Zorya co-localization was enriched in marine samples (Fig. 3c). Similar co-localization trends were observed if only counter-defense genes were considered in the analysis (Supplementary Fig. 3a, Supplementary Data 5).

**Fig. 3.**
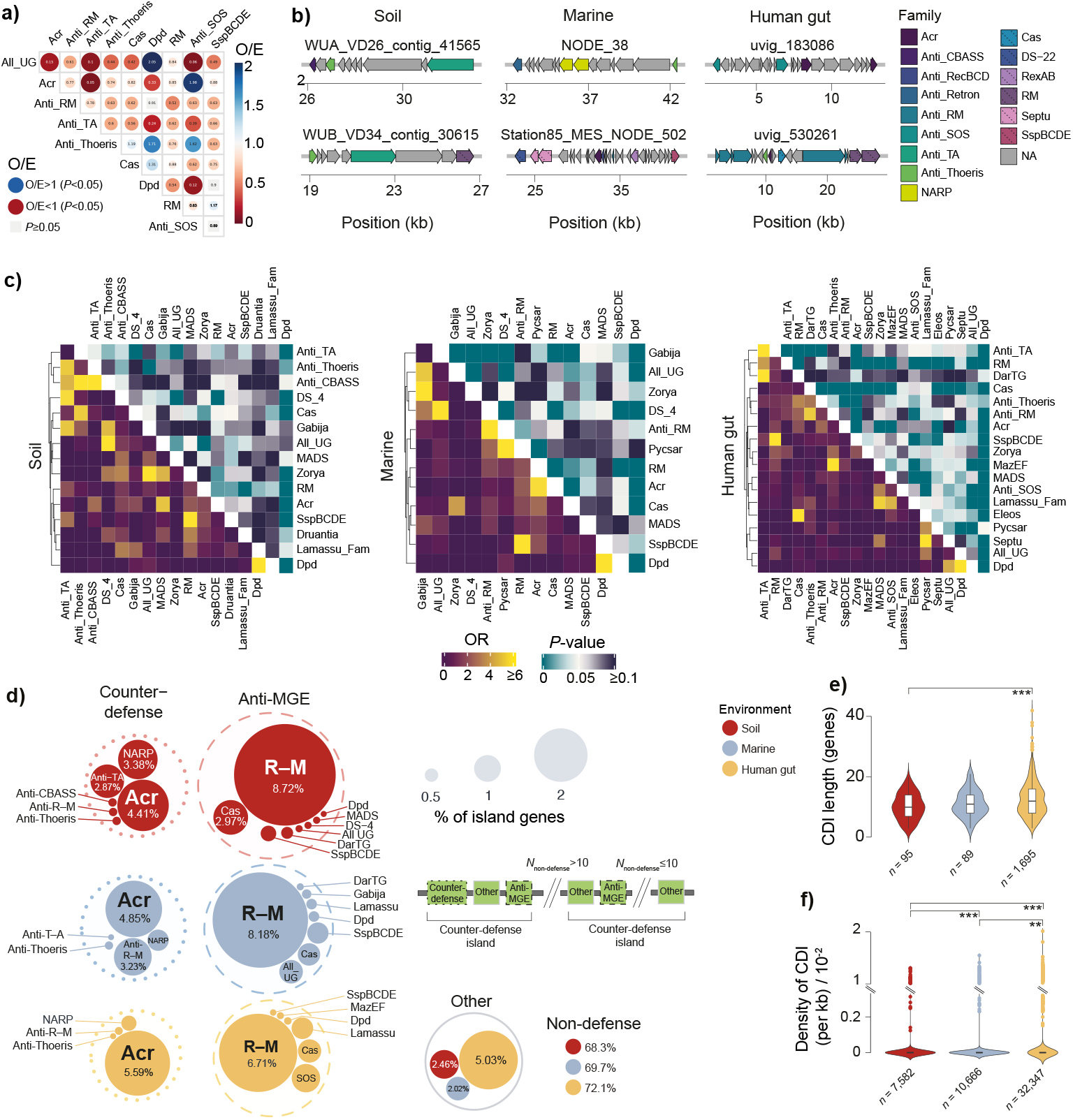
Co-occurrence and co-localization of counter-defense and anti-MGE immune modules across soil, marine, and human gut vOTUs. **a**) Pairwise co-occurrence of presence/absence patterns for the five most abundant counter-defense and anti-MGE families across 36,239 phage genomes. For each family pair, presence/absence was tested for independence using Pearson’s chi-square test, and the ratio of observed to expected (O/E) joint presence was computed, where the expected joint presence was calculated as the product of each family’s marginal presence frequency across the dataset. Circle color indicates the direction of the deviation from the null expectation of independence (red, O/E *<* 1; blue, O/E *>* 1), and both size and color intensity scale with the magnitude of that deviation. The O/E value is given within each symbol. Gray squares indicate non-significant correlations (*P ≥* 0.05, Chi-square test). **b**) Representative examples of co-localizing immune modules across environments. **c)** Odds ratio (OR) of co-localization (bottom heatmaps) and associated two-sided Fisher’s exact test *P* values (upper heatmaps) among anti-MGE defense and counter-defense gene families, restricted to families with a co-localization frequency *≥*1% across soil, marine, and human gut vOTUs. **d**) Relative abundance (%) of gene families within counter-defense islands across environments. Families accounting for less than 0.5% of occurrences were classified as “Other”, whereas all remaining genes were classified as “Non-defense”. Counter-defense islands were defined as genomic regions containing at least three distinct families of counter-defense and/or anti-MGE genes within a neighborhood harboring no more than 10 intervening CDSs. **e**) Counter-defense island (CDI) length distribution (in genes) across environments. **f**) Distribution of CDI densities (per kb, per vOTU). Statistical comparisons were performed using two-sided Wilcoxon rank-sum tests (*^∗∗^P <* 10*^−^*^2^*,^∗∗∗^ P <* 10*^−^*^3^).

We next quantified the extent to which these co-localized genes are organized within counter-defense islands (Methods). Counter-defense and anti-MGE gene clusters colocalized within 10 CDS of one another in 3.7% of phage genomes, with 26.7% co-localizing as immediately adjacent blocks. In total, we identified 99 counter-defense islands in 93 vOTUs from our dataset (Supplementary Data 6, Methods). Although more than 70% of the genes within counter-defense islands could not be assigned a clear immune-related function, approximately 20 major defense-associated families were identified, with R–M, Cas, Acr and NARP being the most abundant (Fig. 3d). These islands displayed similar median sizes across environments (approximately 10 genes), although human-associated samples contained the largest islands of up to 40 genes (Fig. 3e). Counter-defense island density likewise showed comparable median values across environments, but differed in its distribution, with soil and marine samples exhibiting the narrowest ranges and human-associated samples the greatest variability with as much as 3 DIs per vOTU (Fig. 3f). These patterns are consistent with earlier observations of an expanded defensome and increased island content in human gut MAGs relative to those from other biomes ^15^.

Recent observations stemming from Genbank data, indicate that the homing endonuclease SegB facilitates immune evasion in phages by promoting segmental amplification of counter-defense loci containing tRNAs ^46^. In our dataset, we identified 315 instances of *segB*, along with tRNAs in its close vicinity, but contrary to previously observed ^46^, *segB*-vicinal regions exhibited lower median tRNA densities than those observed across the corresponding full genomes (Supplementary Fig. 3b, Supplementary Data 7,8). The exception to this trend consisted in vOTUs belonging to the *Steigviridae* family, which are known to harbor exceptionally high numbers of tRNAs ^47^, and for which we observed the highest values of tRNA denisty around *segB*. This pattern suggests that SegB-mediated counter-defense may be more strongly dependent on phage family context than previously appreciated.

In summary, co-localization patterns point to potential unrecognized epistatic relationships among specific subsets of counter-defense and anti-MGE genes in phage genomes. Approximately 8% of phage immune-related modules are concentrated within counter-defense islands, which display environmental variation and contain a substantial proportion of non-defense genes. Similar to bacterial defense islands, these genomic hubs are likely to shape phage–host antagonistic interactions.

### Biogeography shapes phage counter-defense repertoires and transcriptome

Environmental conditions shape phage diversity, and phages in turn shape the microbial composition of their environments, driving continuous co-evolution. To explore this interplay, we examined how the phage counter-defensome varies with biogeography. Counter-defense systems exhibited marked heterogeneity in both density and family composition across finer ecological sub-habitats in soil and marine environments and across geographic cohorts in the human gut (Fig. 4a, Supplementary Data 9). Soil vOTUs from desert environments displayed particularly high counter-defense densities and anti-SOS modules, whereas volcanic soils were comparatively depleted and dominated by Acr modules. Marine phages maintained lower overall densities with a homogeneous breakdown of counter-defense families across oceanic regions. In contrast, human-gut phages sustained consistently higher densities across countries, with broadly similar frequency spectra interrupted by clear country-specific differences (for example, virtual absence of Acr systems in Italian samples contrasted with their marked over-representation in Tanzanian cohorts). Counter-defense modules, including anti-SOS elements, were more abundant in human gut phages than in marine or soil phages, in line with the higher density and diversity of bacterial defenses, as well as frequent SOS induction in the gut due to genotoxic stressors and high prophage induction rates ^48–50^. The results were qualitatively similar when performed with counter-defense genes (Supplementary Fig. 4a, Supplementary Data 9) or anti-MGE genes (Supplementary Fig. 4b, Supplementary Data 9), with the exception that overall densities were globally more elevated in the latter.

**Fig. 4.**
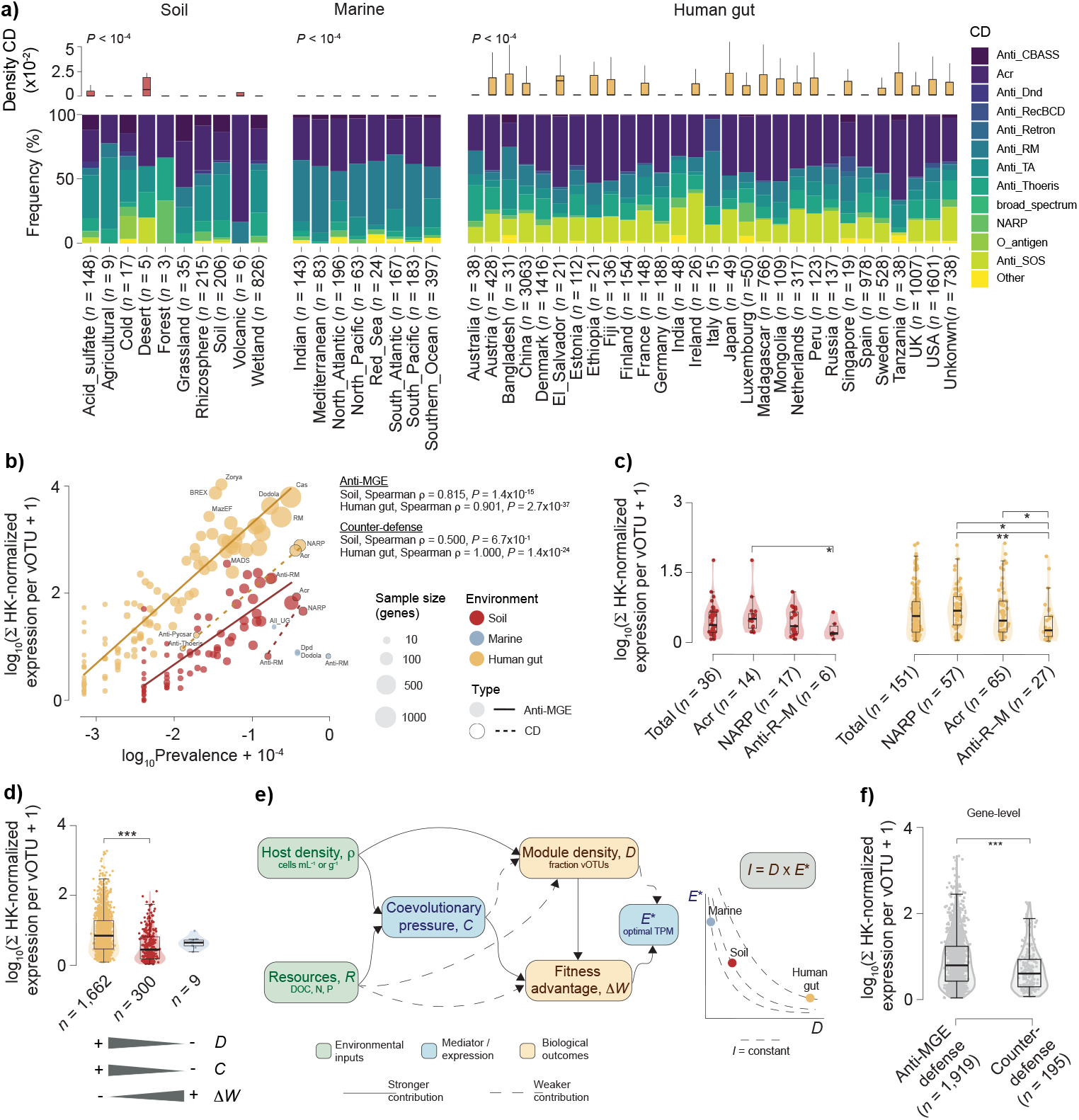
Interplay between phage biogeography, counter-defense repertoires, and their transcriptome. **a)** Density and frequency of counter-defense (CD) systems across distinct ecological (soil, marine) and geographical (human gut) contexts. Boxplots depict the interquartile range (25th - 75th percentiles), with the median indicated by a central black line. Whiskers extend to 1.5x the interquartile range, and outliers were omitted for improved visualization. Statistical significance was assessed using a Kruskal-Wallis test. The number of vOTUs included in each category is provided in parentheses. **b)** Relationship between gene prevalence and sum of housekeeping (HK)-normalized expression per vOTU across anti-MGE defense (solid trend lines) and counter-defense (dashed trend lines) genes in soil (red), human gut (yellow), and marine (blue) samples. Each point represents one immune family sized by the number of contributing genes. Spearman correlation coefficients and *P* values are shown for each immune class and environment. Marine was excluded from correlation testing owing to small sample size. **c**) Summed HK-normalized expression per vOTU of counter-defense genes carried by soil (red) and human gut (yellow) vOTUs, shown for all counter-defense-carrying vOTUs pooled (Total) and split by family. *n* corresponds to the number of vOTUs per category, and only those with *n ≥* 5 are shown. **d**) Same expression metric as in (b, c), pooled across all counter-defense + anti-MGE genes and compared across environments. *n* corresponds to the number of vOTUs per category. Beneath the plot, schematic gradients summarize how immune module density (*D*), coevolutionary pressure (C), and per-module fitness advantage (Δ*W*) are expected to vary across environments. **e**) Proposed model linking environmental inputs to realized immune investment. Host density (*ρ*) and resource availability (*R*) jointly shape co-evolutionary pressure (*C*), which in turn shapes the fitness advantage (Δ*W*) conferred by carrying a given module. Host density additionally shapes immune module density (*D*, fraction of vOTUs carrying a given module) directly, independent of co-evolutionary pressure, with *C* exerting a minor, secondary influence on *D* over longer timescales. *D* feeds directly into Δ*W* (as a module becomes more prevalent, its marginal fitness benefit erodes through negative frequency-dependence). Δ*W* is the primary driver of each module’s optimal expression level (*E^∗^*). *D* contributes a smaller, secondary effect, in line with a redundancy/free-riding dynamic. Dashed arrows denote weaker, secondary contributions alongside the primary (solid) pathways above. Total realized immune investment is modeled as *I* = *D* × *E^∗^*. The graph on the right represents optimal expression (*E^∗^*) against module density (*D*) for the three environments. Realized total investment (*I*) is highest in human gut and comparably low in soil and marine. Human gut and marine sit at opposite, constrained extremes of the *D*–*E^∗^* trade-off. In the gut, *D* saturation (via HGT) reduces the fitness advantage of the module, pulling *E^∗^* down even as *D* itself stays high. In marine, a genome-streamlining selection filter caps *D* directly, even though per-carrier *E^∗^* remains comparatively high. Soil is unconstrained by either extreme, but its overall inputs (*ρ*, *C*) are modest, so it remains at low *D* and low-to-moderate *E^∗^*. **f**) Summed HK-normalized expression per vOTU of immune genes split by anti-MGE or counter-defense function. *n* corresponds to the number of vOTUs per category. Statistical comparisons were performed using two-sided Wilcoxon rank-sum tests (*^∗^P <* 0.05*,^∗∗^ P <* 10*^−^*^2^*,^∗∗∗^ P <* 10*^−^*^3^)

Having analyzed the impact of biogeography, we next asked which counter-defense and anti-MGE families were transcriptionally active in viral populations, and at what magnitude. To this end, we mapped metatranscriptomic reads from soil, human gut, and marine viromes onto their cognate vOTUs, and related summed per-vOTU transcript abundance for each family to that gene or system’s prevalence within the corresponding metagenomic population. This analysis revealed that counter-defense and anti-MGE immune modules are transcribed highly unevenly, with transcriptional level tightly coupled to how common a given gene / system is at the population level. Summed per-vOTU expression correlated positively with gene prevalence in both soil and human gut metatranscriptomes (Fig. 4b, Supplementary Data 10). Marine metatranscriptomes, by contrast, were comparatively depleted in immune modules, and only a handful of genes passed detection thresholds (see Methods), precluding a statistically robust correlation. Genes from widely distributed families (Cas, R–M, NARP, Acr, Anti-R–M) mapped at the high-prevalence, high-expression end of this relationship in both environments, while narrowly distributed genes were both rare and weakly transcribed. When comparing the most prevalent counter-defense families in each environment, anti-R–M genes were significantly more weakly expressed than Acr and NARP in both soil and human gut vOTUs (Fig. 4c, Supplementary Data 11). This might be explained by the fact that R–M cleaves unmodified phage DNA shortly after cell entry, and anti-R–M genes are correspondingly restricted to a brief, immediate burst at the point of genome entry, a narrow window that a cross-sectional sample is less likely to catch. Instead, CRISPR interference and *NAD*^+^ depletion unfold over longer and/or later windows of infection ^51–53^, so Acr and NARP may remain transcriptionally active across a substantially larger share of the infection cycle, making them proportionally easier to detect in cross-sectional data. At the community level, total per-vOTU immune-related expression differed sharply between environments, with human gut vOTUs showing significantly higher expression than soil vOTUs, and marine vOTU data being less abundant but falling within the same low range as soil (Fig. 4d, Supplementary Data 12). This trend correlates with lower host density, co-evolutionary pressure, and module density in marine environments relative to the human gut. Yet this same rarity is expected to confer, via negative frequency-dependence, a higher fitness advantage on the few marine phages that carry a given module. These observations can be framed with a simple resource–density model (Fig. 4e). Under such scenario, in densely populated ecosystems like the human gut, elevated viral-host encounter rates typically drive extensive neutral HGT, leading to a ubiquitous prevalence of immune modules across vOTUs. However, this genetic saturation is expected to diminish the marginal fitness advantage of the trait during viral-viral and viral-host competition, as a module carried by a large number of genomes ceases to confer a competitive edge. To offset the metabolic burden of transcription, selection will likely drive optimal expression to basal levels. Conversely, in less populated marine environments, module prevalence is expected to decrease under stringent selection conditions and reduced contact times, but this same decrease will likely maximize the competitive payoff for the few phages possessing these systems, driving localized expression to its theoretical peak. In soil, by contrast, neither constraint dominates, and module prevalence and per-carrier expression both settle at intermediate values. This release from either extreme, however, does not seem to translate into the highest total investment, because soil’s underlying host density and co-evolutionary pressure are themselves comparatively modest, and its realized immune investment remains comparable to that of marine communities. Realized investment is instead highest in the human gut, where the scale of host density-driven module acquisition outweighs the accompanying reduction in per-carrier expression. Similar trends were obtained when performing the analysis at the system level (Supplementary Figs. 5a,b, Supplementary Data 13) and when restricting it to anti-MGE modules alone (Supplementary Fig. 5c,d, Supplementary Data 14). Globally, anti-MGE genes showed significantly higher expression than counter-defense genes (Fig. 4f). As noted above, this asymmetry may reflect a genuine biological difference in transcriptional investment between the two module classes, or the current annotation bias between them. Regardless, system-level resolution revealed that phage-encoded CRISPR/Cas systems dominated anti-MGE expression in the human gut, whereas R–M systems were the most actively expressed in soil (Supplementary Fig. 5d).

Analysis of prophage regions within bacterial MAGs from the same communities revealed an environmental pattern distinct from that observed at the vOTU level, with anti-MGE defense expression highest in marine samples, intermediate in human gut samples, and lowest in soil. Genes pertaining to R–M, Cas, and Dodola were the most abundantly expressed (Supplementary Figs. 6a-c, Supplementary Data 15), whereas only R–M remained when performing the analysis at the system level (Supplementary Fig. 6d, Supplementary Data 15). Expression of these prophage-encoded defense genes and systems is consistent with roles in methylome maintenance, protection against competing MGEs, and persistence of prophage–host associations.

Collectively, these patterns demonstrate that phage counter-defense repertoires are finely tuned to local ecological and geographic contexts, shaped by host defenses, environmental stressors, and evolutionary pressures. They also support a model in which transcriptional investment, and the specific systems recruited to deliver it, scale primarily with host density and the resulting intensity of host-phage arms races.

### Immune-related modules are absent in virophages but present in NCLDVs

The widespread occurrence of anti-MGE defenses in bacteria and counter-defenses in bacteriophages, together with evidence of HGT involving bacteria, phages, virophages, and NCLDVs, suggests that immune-related genes may also be present in virophages and NCLDVs. To explore this hypothesis, we systematically surveyed the presence and distribution of bacterial anti-MGE defense genes and phage counter-defense genes in virophage and NCLDV genomes (Supplementary Fig. 7a, Supplementary Data 16). We detected no significant homology between virophage genes and any characterized immune modules from prokaryotes or phages. Given the highly compact nature of virophage genomes and their obligate dependence on the replication machinery of NCLDVs, virophages are likely to rely on minimal, structurally simple counter-defense strategies. Furthermore, the selective pressures imposed by giant virus-encoded defense systems such as MIMIVIRE, are expected to drive accelerated sequence evolution, recombination, or other genomic innovations, that enable virophages to evade targeting while maintaining parasitic compatibility with their hosts. On the other hand, across 5,284 NCLDV genomes, we identified only three instances of MIMIVIRE, and a limited number of phage counter-defense systems and genes (126 and 776 respectively) (Fig. 5a, Supplementary Data 17). In contrast, a diverse repertoire of bacterial anti-MGE defense genes was observed, unevenly distributed across the dataset (Supplementary Data 17). Some families, notably CRISPR/Cas and R–M, were present in a substantial fraction of genomes (*>* 39%), whereas others were confined to a limited number of viral lineages. To assess whether these genes occur as solitary elements or as components of larger systems, we examined their genomic organization. Solitary genes were the most abundant, particularly DNA MTases, while multi-gene configurations were comparatively rare. An exception was the presence of digenic arrangements, consistent with the presence of R–M modules (Fig. 5b). Among MTases, Type II genes dominated, whereas Type I were less frequently observed (Fig. 5c). We also identified loci in which multiple defense genes co-localize, although these were less frequent than islands reported in bacterial or phage genomes (Supplementary Fig. 7b, Supplementary Data 18). When comparing NCLDV-encoded defense-gene abundance across biomes, marine was the most NCLDV-rich environment overall, and also the one in which these viruses harbor the highest density of defense genes (Fig. 5d). Within marine samples specifically, no significant differences were observed in defense-gene family composition across different oceans (Supplementary Fig. 7c). When splitting per defense and NCLDV families, the most abundant viral families (*Mimiviridae* and *Phycodnaviridae*) exhibited a statistically significant enrichment of multiple defense genes, whereas a small subset of families, such as Dodola, Lamassu, and PsyrTA, were enriched across most viral families (Fig. 5e). Concomitantly, the least abundant viral families, were largely devoid of most known defense genes, with particular emphasis to *Asfarviridae*, in which only Dnd was detected.

**Fig. 5.**
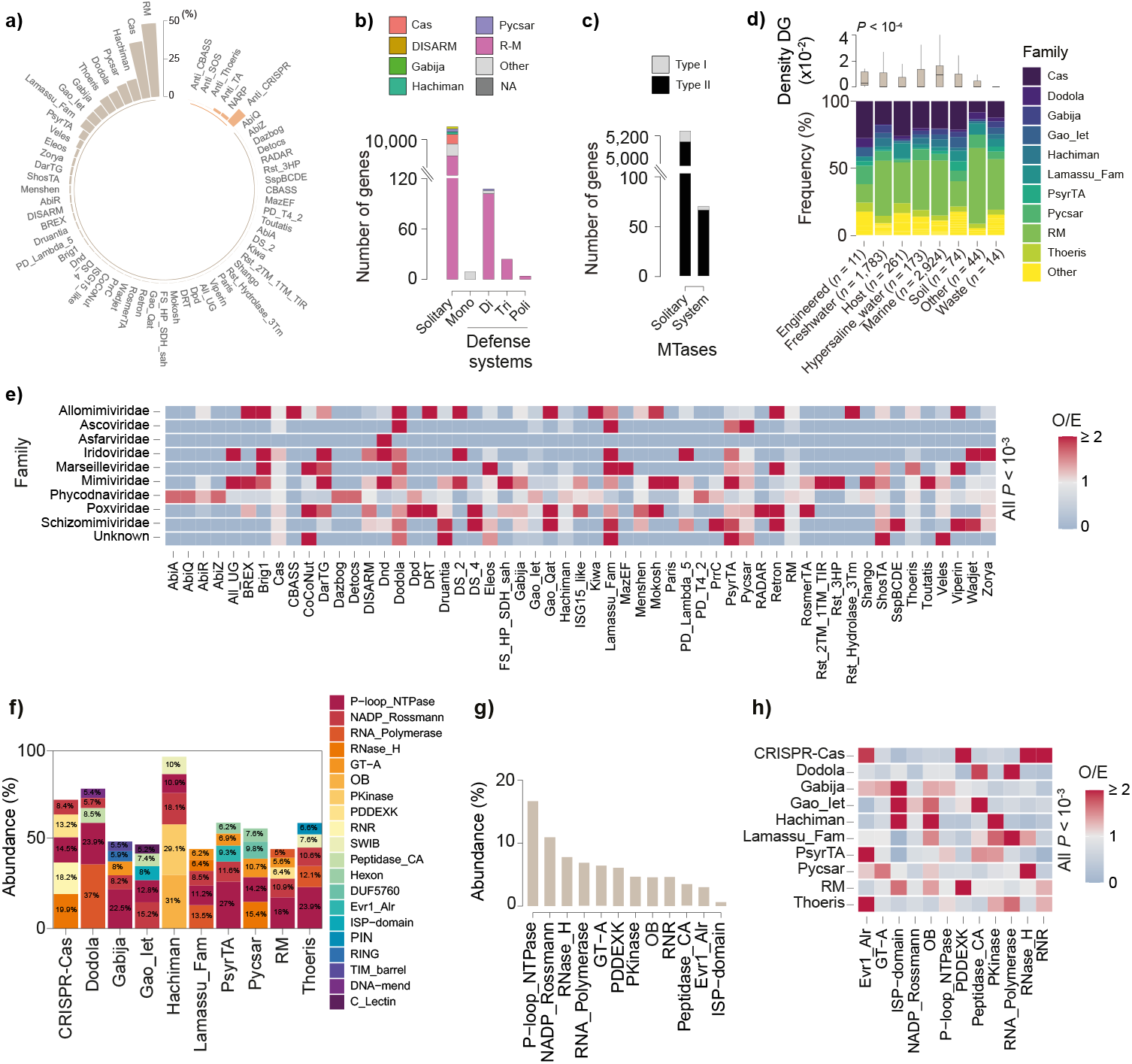
Bacterial anti-MGE defense genes are abundant in NCLDVs. **a**) Fraction of NCLDVs encoding each identified anti-MGE defense family (beige) compared to counter-defense families (light orange). **b**) Distribution of defense genes (DG) according to genomic organization (solitary or arranged within mono-, di-, tri-, or polygenic defense systems). **c**) Classification of DNA MTase genes as either solitary or belonging to complete R–M systems. **d**) Density and frequency of anti-MGE genes in NCLDVs recovered from distinct environments. Boxplots depict the interquartile range (25th - 75th percentiles), with the median indicated by a central black line. Whiskers extend to 1.5x the interquartile range, and outliers were omitted for improved visualization. Statistical significance was assessed using a Kruskal-Wallis test. The number of NCLDV genomes included in each category is provided in parentheses. **d)** Heatmap of observed-to-expected (O/E) ratios for distinct defense families across major NCLDV families. Expected values were calculated by multiplying the total number of genes within each defense family by the overall proportion of all defense-associated genes assigned to each viral family across the full dataset. **f**) Most abundant protein domain clans found within a ± 4 CDS window surrounding the ten most abundant NCLDV defense gene families (top 5 clans per family shown). **g**) Rank-ordered abundance of these domain clans within the same ± 4 CDS genomic radius. **h**) Heatmap of observed- to-expected (O/E) ratios for domain clans across major defense gene families. Expected values were calculated by multiplying the total number of domains within each clan by the overall proportion of all domains assigned to each defense family across the full dataset.

To further characterize the genomic context of immune-related genes in NCLDVs, we analyzed protein domain composition within a ±4 coding sequence window around known defense gene anchors. This analysis revealed significant enrichment of several domain clans, many of which have established or putative roles in antiviral defense (Figs. 5f-g, Supplementary Data 19). Examples include: *i*) Cas4-like domains, a hallmark component of the MIMIVIRE system; *ii*) members of the PD-(D/E)XK nuclease superfamily ^54,55^; *iii*) RNase H and RNase H-like domains ^56^; *iv*) OB-fold kinases ^57^; *v*) RING finger domains, which function as putative E3 ubiquitin ligases ^58^; *vi*) ribonucleotide reductases (RNRs) ^59^; and *vii*) GT-A fold glycosyltransferases ^60^. Observed-to-expected enrichment analysis revealed non-random co-occurrence between defense-family homologs and specific accessory domain clans (Fig. 5h). As these families are assigned by homology to defense systems characterized in bacteria and archaea, whether the associated domains reflect genuine functional coordination, shared horizontal acquisition, or broader features of NCLDV genome organization remains to be determined.

Overall, these results reveal a clear contrast between virophages and NCLDVs in the presence of immune-related genes. While no immune modules were detected in virophages, NCLDVs encode a diverse set of genes homologous to bacterial anti-MGE defense systems, unevenly distributed across viral lineages and predominantly present as solitary loci, with occasional multi-gene configurations. These loci show non-random genomic association with putative defense-associated accessory domains, suggesting that NCLDVs have evolved a mosaic defensive landscape either reflecting functional innovation, differential retention, or recurrent horizontal acquisition.

## Discussion

In this study, we present a comprehensive analysis of the abundance and diversity of phage-encoded counter-defensomes across three representative biomes: soil, marine environments, and the human gut. This work complements previous analyses of bacterial MAG defensomes ^15^, enabling a parallel, biome-resolved comparison of defense and counter-defense strategies across both hosts and their viruses. Consistent with prior studies on Refseq genomes ^20^, we found predominance of anti-R–M and Acr proteins, presumably because their targets are also the most abundant in bacterial populations. Although counter-defense genes were detected across a broad diversity of vOTUs, their distribution was skewed toward phages predicted to infect *Gammaproteobacteria*. This apparent enrichment likely reflects historical discovery bias, as early characterization of counter-defense systems relied heavily on model Gammaproteobacterial hosts and their associated phages. Furthermore, detection biases may contribute to these patterns, as taxa overrepresented in reference databases such as RefSeq are not necessarily the most abundant in natural environments, whereas ecologically dominant but under-sequenced organisms may be systematically underrepresented in current annotations of counter-defense genes. Challenges remain in the detection of distantly related counter-defense proteins, largely due to their small size and extensive sequence divergence. Recent advances in protein structure prediction and comparative structural analyses are helping to overcome these limitations, enabling a shift toward structure-guided discovery of novel counter-defense systems. This approach has already proven effective in identifying new Acr proteins in bacteriophages ^61^. In any case, the diversity of counter defense systems we detected is consistent with recent simulations predicting the number of immune modules sustained in coexisting ecosystems ^62,63^.

Counter-defense modules were found to be globally more abundant in human gut phages than in marine or soil phages (Fig. 2a). This enrichment likely stems from the elevated density and diversity of bacterial defense systems in the gut ^15^, which impose stronger and more continuous selective pressure on phages despite the characteristically low VPR and predominance of temperate lifestyles. Constant exposure to a high density and diversity of defense systems is expected to drive rapid phage adaptation through bursts of acquisition of compatible counter-defense systems or the expansion of existing ones ^64^, with similar adaptive dynamics likely to occur in other defense-rich environments, including the urethra, vagina, oral cavity, and skin ^65^. We also observed an enrichment of anti-SOS response modules in gut-derived phages. This pattern is consistent with the higher frequency of SOS response induction in the human gut microbiome than in marine or soil habitats, likely driven by persistent genotoxic stressors such as host-derived reactive oxygen and nitrogen species, microbial secondary metabolites (*e.g.*, colibactin), bile salts, and frequent prophage induction linked to high lysogeny rates ^48–50^.

Similarly to the bacterial defensome ^15^, the phage counter-defensome also displayed marked biogeographic variation. In particular, desert environments exhibited a higher density of counter-defense genes, whereas volcanic environments were characterized by comparatively lower diversity. Drought conditions were previously shown to nearly double negative prokaryote–phage interactions, coinciding with an increased abundance and diversity of bacterial antiphage defense systems ^66^. In the particular case of microbial communities in hyper arid desert soils, they experience episodic hydration alongside persistent exposure to desiccation, osmotic, and UV-induced oxidative stresses. These environments are further characterized by a high prevalence of lysogeny and low extracellular viral abundances ^67^. Such conditions may explain the selection for vOTUs enriched in anti-SOS systems (Fig. 4a), potentially as part of an adaptive strategy to suppress host stress–induced prophage induction. In the case of volcanic soils, SO_2_ exposure creates a strongly filtered habitat with the most affected deposits being acidic, carbon- and nitrogen-poor, and dominated by a small set of pioneer autotrophs. In these rapidly assembling bacterial communities of newly formed volcanic soils, phage-encoded counter-defense systems are expected to exhibit limited diversity, reflecting the transient and taxonomically dynamic host landscape ^68^. Instead of maintaining a wide repertoire of highly specialized modules tailored to individual hosts, phages in these environments are likely to rely on broadly effective strategies, which helps explain the observed predominance of anti-R–M and Acr mechanisms (Fig. **??**). As microbial communities stabilize during ecological succession, the increasing diversity and persistence of host interactions are expected to drive the expansion and diversification of the phage counter-defensome.

We found extensive co-localization between the phage counter-defensome and the anti-MGE defensome, as well as general evidence for the existence of counter-defense islands. This pattern supports the view that MGEs are major drivers of Red Queen dynamics by promoting the rapid dissemination and co-accumulation of both defense and their counter-defense mechanisms within microbial populations. We also provided, the first snapshot of the metatranscriptome landscape of immune modules for phage populations across diverse biomes. To date, no community metatranscriptomic study has syste matically quantified the expression of known anti-MGE defense and counter-defense genes across natural microbiomes. Instead, most ecological studies infer immune repertoires from gene presence in metagenomes, whereas expression analyses have largely been restricted to laboratory infection studies ^69–71^. For example, recent work in *Escherichia coli* ^72^ showed that bacterial anti-MGE transcription varies with environmental conditions, growth stage, and genetic background, even without experimentally imposed phage infection. These findings complement our biome-resolved metatranscriptomic observations and suggest that variation in viral immune-related expression may reflect physiological and regulatory differences alongside module prevalence and host–virus co-evolutionary pressures.

We caution, however, that assigning expression signals to viral or bacterial origins is not unequivocal. Since read assignment relies on competitive pseudoalignment, transcripts originating from conserved or recently horizontally transferred loci may align equally well to homologs encoded in both viral and bacterial genomes and are therefore assigned probabilistically according to their relative transcript abundance. Yet, since bacterial transcripts typically dominate community metatranscriptomes, ambiguous reads are expected to be preferentially assigned to the bacterial compartment, making our estimates of phage defense and counter-defense expression conservative.

The diversification of immune systems is often facilitated by their modular organization, which allows frequent recruitment and exchange of defensive modules. In recent years, our understanding of the anti-MGE defensome has expanded dramatically, largely driven by the ‘guilty by association’ principle, *i.e.*, the observation that multiple immunity-related modules frequently cluster within specific genomic loci in bacteria ^73^. In this study, we extended this concept to NCLDVs, querying domains preferentially co-localized with known immune modules. While the functional significance of these co-localizations remains uncertain, their enrichment in defense-associated domains is consistent with a role in coordinated antiviral activity. Our observations in NCLDVs align with recent findings using machine-learning frameworks, which identified PD-(D/E)XK nucleases, P-loop NTPases, and PIN domains among the most prevalent features in newly predicted defense modules in bacteria ^61^. Also, our identification of numerous Type II MTases helps explain the methylome diversity of giant viruses ^37^. Moreover, the predominance of solitary enzymes suggests that at least some may play functional roles in gene expression regulation. It remains to be determined to what extent bacterial MTases carried by NCLDVs are transferred to eukaryotic hosts, how they evolve within NCLDV genomes, and whether, similarly to bacteria, viruses with comparable methylation repertoires promote genetic exchange ^74,75^. Hence, our study provides a biome-resolved view of phage anti-MGE and counter-defense repertoires, extended to dsDNA viruses. These insights refine our understanding of host–virus interactions across ecosystems and establish a framework for the discovery of novel immunity-related modules across the virosphere.

## Methods

### Data

In this study, we analyzed a large dataset of 50,595 publicly available phage vOTUs from soil ^76–79^, marine ^38,80–82^, and human gut environments ^40^. These vOTUs were filtered using PhaBOX^83^ v2.1.12 (default parameters) to retain only viral contigs, further screened for genome quality using CheckV ^84^ v1.0.3 (end_to_end mode, completeness *≥*90%, contamination *≤*5%), and dereplicated using dRep ^85^ v3.6.2 (options: -pa 1, -l 18000) to remove redundant sequences and retain only high-quality contigs. vOTU annotation was performed with Prokka ^86^ v1.14.5 (default parameters). In parallel, we analyzed dedicated datasets of 1,682 virophage ^87^ and 5,284 NCLDV genomes ^88,89^. Contigs were dereplicated using dRep (-pa 1), with an additional minimum length threshold of 10 kb applied to virophage sequences. For the detection and quantification of immune gene expression, 19 publicly available metatranscriptomic datasets were used, spanning marine, soil (^90–92^), and human gut environments (^93–95^) (Supplementary Data 10). These studies also provided corresponding metagenomic data, enabling: i) the recovery of vOTUs from CheckV-filtered contigs *≥* 1 kb, and ii) the recovery of bacterial metagenome-assembled genomes (MAGs) quality-assessed with CheckM v1.2.3 (completeness *≥* 90%, contamination *≤* 10%). Both viral and bacterial genome catalogs were dereplicated with dRep at a 95% average nucleotide identity (ANI) threshold. Gene prediction was performed with Prodigal ^96^ v2.6.3.1. High-confidence prophage regions were identified from bacterial MAGs with VirSorter2^97^ (score *≥* 0.9), and only full viral sequences *≥* 1 kb in length were retained for further analysis.

### Taxonomic assignment and viral morphology

Taxonomic assignment of phage vOTUs was performed using a combination of PhaBOX (default parameters), Kaiju ^98^ v1.7.3 (default parameters), and vConTACT3^99^ v3.1.6 (options: –nucleotide –db-version 230). vOTUs sharing *≥*20% of their homologous proteins were considered to belong to the same viral family. Unclassified sequences were further assigned at the family level using VPF-class ^100^ (membership ratio *>*0.5, confidence score *>*0.5), retaining only the highest-confidence prediction per genome. Host taxonomy was predicted using PhaBOX and iPHoP ^101^ v1.4.1 (default parameters). Viral morphology was inferred from the functional annotation of structural proteins by comparing predicted proteins against the RefSeq viral database using MMseqs2^102^ and the non-redundant database using DIAMOND ^103^ v2.0.15, with assignments based on the detection of characteristic marker proteins (major capsid protein, portal protein, and tail proteins for head-tailed viruses, filamentous capsid proteins for rod-shaped viruses, and double jelly-roll capsid proteins for icosahedral viruses). For vOTUs that remained unclassified, morphological information was further refined by propagation within gene-sharing clusters defined by vConTACT3. Although *Archaea* are also infected by viruses, there is ongoing debate about whether these viruses should be classified as phages. In this study, archaea-infecting viruses were considered phages, despite the fact that most viruses considered in this study typically infect bacteria. For virophage and NCLDVs, taxonomic assignments were verified using PhaBOX and Kaiju, retaining only sequences classified as *Lavidaviridae* for virophages and *Nucleocytoviricota* for giant viruses.

### Identification of anti-MGE defense and counter-defense genes / systems and islands

Anti-MGE defense genes and systems were identified in vOTUs using DefenseFinder ^14^ v2.0.0 (option: –preserve-raw). Counter-defense genes and systems were identified by combining DefenseFinder anti-defense mode (options: –antidefensefinder-only –preserve-raw) with the dbAPIS framework ^104^. For dbAPIS-based detection, predicted proteins were screened using both profile-based searches with HMMER v3.4 (http://hmmer.janelia.org/) (hmmscan) and sequence similarity searches with DIAMOND ^103^ v2.0.15 (blastp), and annotations were integrated using the dbAPIS parsing workflow. Hits were filtered to retain only alignments with a minimum coverage of 80%, and for each gene only the best-supported assignment was kept. Finally, overlapping annotations between defense and counter-defense catalogs were removed at both the gene and system levels. MIMIVIRE systems were identified through a BLASTp search between the protein sequences R349 (YP_003986852.1), R350 (YP_003986853.1), and R354 (YP_003986857.1) from *Acanthamoeba polyphaga* mimivirus (NC_014649.1) and the full NCLDV proteome dataset. Hits were filtered at a minimum of 80% identity and 80% query coverage. To assess co-localization between anti-MGE / counter-defense families, we computed odds ratios and associated Fisher’s exact test *P* values. For this purpose, counter-defense genes were ordered along each contig and co-localization was evaluated using a bidirectional sliding window in which each annotated gene was paired with up to the next five annotated genes on the same contig. Because genes belonging to the same predicted complete system are inherently clustered, gene pairs assigned to the same system were deliberately excluded to avoid inflating co-localization frequencies. Counter-defense islands were defined as clusters of annotated genes separated by ten CDS or less, and only clusters containing at least three counter-defense / anti-MGE genes were retained. For islands containing only counter-defense genes, the same identification and filtering strategy was applied.

### Metatranscriptomic analyses of immune genes

Raw paired-end RNA-seq reads were quality- and adapter-trimmed with fastp ^105^ v0.24.0 (options: -q 25, -e 25, -u 20, –length_required 50, –detect_adapter_for_pe). Prokaryotic and eukaryotic non-protein-coding rRNA sequences were then removed from the trimmed paired reads with SortMeRNA ^106^ v4.3.4 (options: –paired_in, –out2, –num_alignments 1). Read pairing was restored after filtering with repair.sh from BBTools ^107^ v39.19, and pseudoaligned against a combined Kallisto ^108^ v0.52.0 index built from the pooled CDS catalogs of the dereplicated bacterial MAGs and viral vOTUs using kallisto quant (options: paired-end mode, –pseudobam, –fr-stranded, –rf-stranded, or unstranded). Pseudoalignment BAM files were coordinate-sorted and indexed with SAMtools ^109^ v1.21, and per-base read depth was computed with sam-tools depth -aa. For each CDS, mapped reads and effective length were taken from Kallisto’s output, and reads per kilobase (RPK) were calculated as 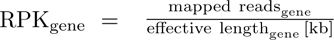 Breadth of coverage was computed from the samtools depth output as the proportion of each gene’s length covered by *≥* 1 read. A gene was considered reliably supported and retained for downstream analysis if it had *≥* 5 mapped reads and *≥* 50% breadth of coverage. Each immune gene’s RPK was normalized to a community-internal housekeeping-gene baseline composed of bacterial *gyrA* and *rpoB* as 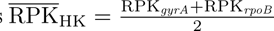. Every supported gene’s housekeeping (HK)-normalized expression was then calculated as 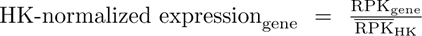. For each vOTU, HK-normalized expression values of all supported defense- or counter-defense-annotated genes were summed to obtain the Σ HK-normalized expression per vOTU.

### Gene neighborhood analyses

To explore the genomic context of defense genes, we performed a gene neighborhood analysis on giant virus contigs. For each defense gene predicted by DefenseFinder, neighboring genes within a sliding window of four coding sequences upstream and downstream on the same contig were retrieved. Defense genes themselves were subsequently excluded from this neighborhood dataset to focus on putative novel defense-associated genes. The resulting protein sequences were functionally annotated using InterProScan ^110^ v5.75.106.0, and only domains assigned to Pfam entries were retained (e-value *≤* 10*^−^*^5^). When available, Pfam clans were used to group related domains into higher-level functional categories.

For *segB* neighborhood analyses, we followed a procedure similar to that described previously ^46^. Briefly, homologs of the T6 phage SegB sequence were identified using BLASTp with an E-value cutoff of 10*^−^*^5^, a minimum sequence identity of 30%, and a minimum coverage of 80%. The resulting sequences were clustered using MMseqs2 v17.b804f easy-cluster with 95% sequence identity and 95% coverage thresholds. The representative sequences from each cluster were then aligned using MAFFT v7.526 with default settings, and an HMM profile was constructed using hmmbuild. For each *segB* homolog, sequences spanning 1 kb upstream and 1 kb downstream were retrieved in FASTA format. Bacterial tRNA genes within these regions were identified using tRNAscan-SE v2.0.12 with default settings. To estimate the confidence interval for the difference in median tRNA counts between *segB*-adjacent regions and the genome-wide background, we performed nonparametric bootstrap resampling with 10,000 iterations. In each iteration, samples were drawn independently with replacement from each group, and the difference between the resulting medians was calculated. The 95% confidence interval was defined by the 2.5th and 97.5th percentiles of the resulting distribution of median differences. To compare tRNA density between *segB*-adjacent regions and the genome-wide background, we used the Wilcoxon signed-rank test. Only genomes for which both *segB*-adjacent tRNA counts and genome-wide tRNA counts were available were included in the analysis.

### Statistical and graphical analyses of data

Statistical and graphical analysis of the results was conducted using R v4.3.3. Sankey diagrams were built with the R package networkD3.

## Code availability

Wrapper scripts supporting all key analyses of this work are publicly available at https://github.com/oliveira-lab/Counter-Defensome.

## Author contributions

P.H.O. supervised the project. A.B., L.S.S., I.G.M.G., L.D. and P.H.O. designed the computational methods. A.B., L.S.S., I.G.M.G., and L.D. performed most of the computational analyses and developed most of the scripts that support the analyses. A.B., L.S.S., I.G.M.G., L.D., N.W., P.W., and P.H.O. analyzed the data. A.B. and P.H.O. wrote the manuscript with additional information inputs from other co-authors.

## Declaration of interest

The authors declare no competing interests.

## Acknowledgments

We thank Eduardo P.C. Rocha (Institut Pasteur, Paris) for his critical reading of the manuscript. This work was supported by the Genoscope, the Commissariat à l’Énergie Atomique et aux Énergies Alternatives (CEA), France Génomique (ANR-10-INBS-09–08), and the Fondation pour la Recherche Médicale (grant number FDT202504020397) to A.B.

## Supplementary Information

**Suppl. Fig. 1.**
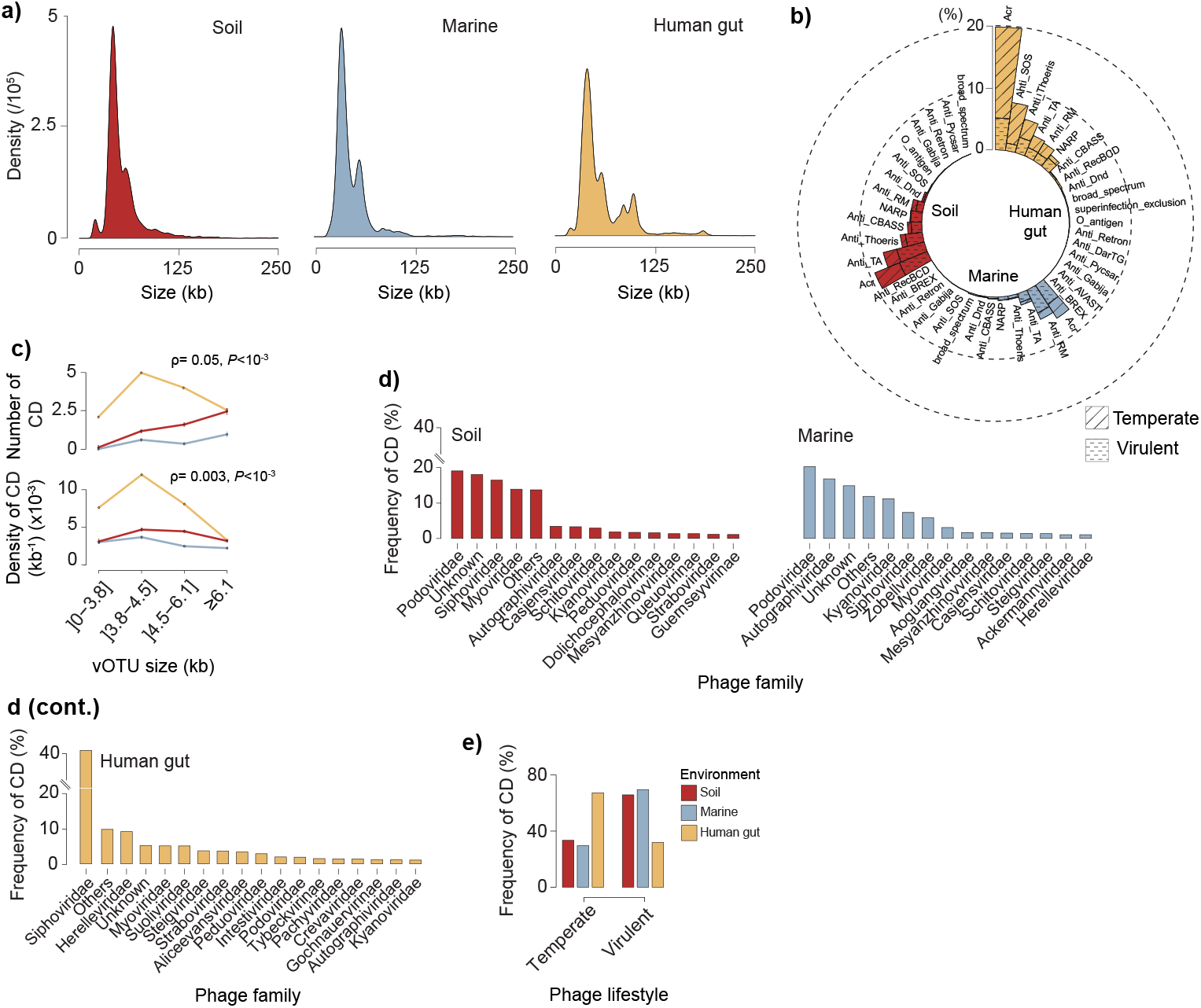
Phage size distribution and ecological partitioning of viral counter-defenses. **a**) Size distribution patterns of the phage dataset. **b**) Distribution of CD genes across phage lifestyles and environments, expressed as percentages. **c**) Changes in the number and density of counter-defense (CD) systems per vOTU (per genome and per kb) as a function of vOTU genome size (kb) across biomes. Error bars indicate standard deviations of the mean. Correlations were assessed by a two-sided Spearman’s rank test. **d**) Distribution of CD systems across phage families and environments, shown as percentages. **e**) Proportion of soil, marine, and human gut vOTUs encoding each CD gene family.

**Suppl. Fig. 2.**
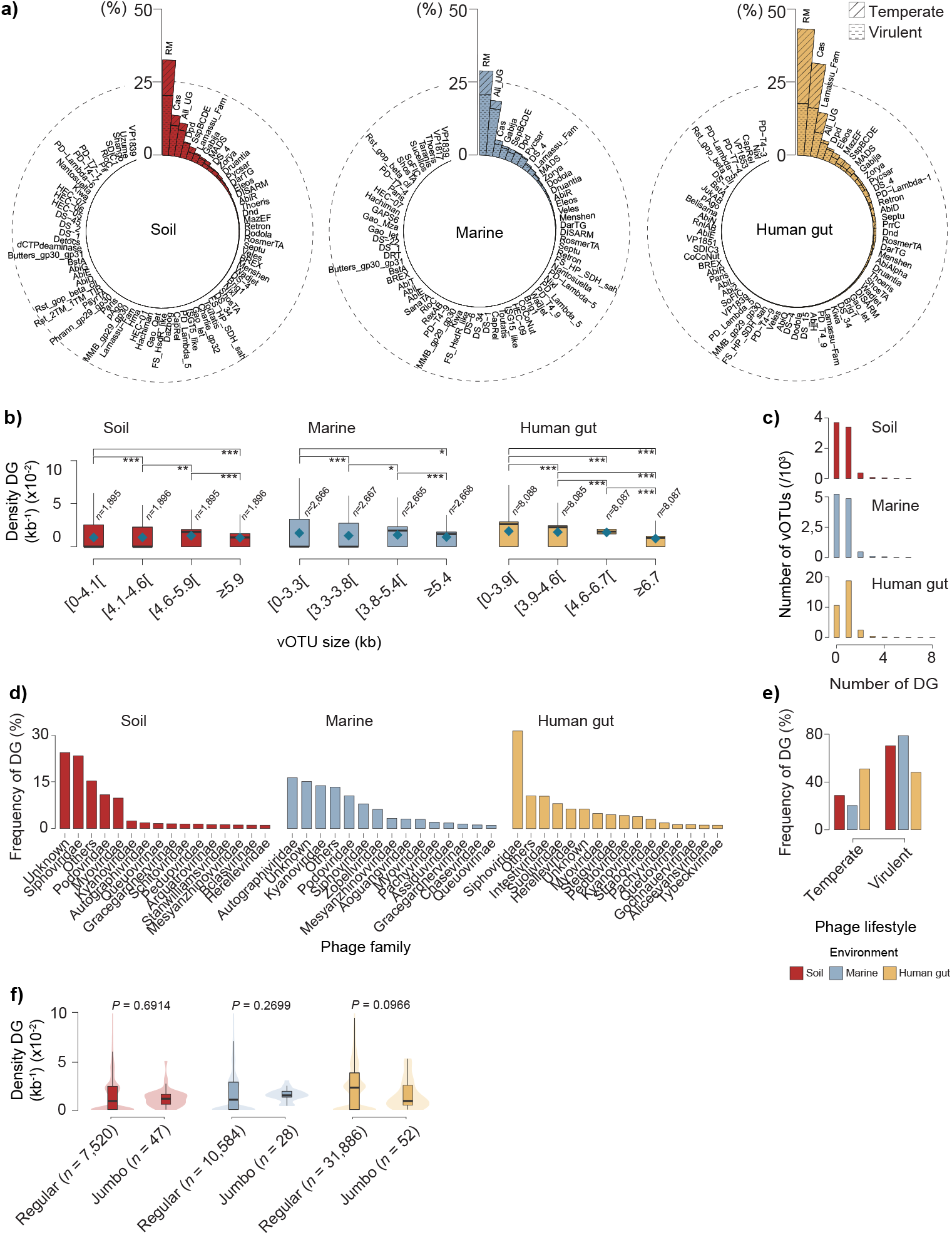
Distribution of anti-MGE defense genes (DG) across viral genomes and environments. **a**) Percentage of soil, marine and human gut vOTUs encoding each family of anti-MGE DG. Data are shown for both temperate and virulent phages. **b**) Relationship between DG density (per kb) and vOTU genome size (kb) across environments. Statistical comparisons were performed using two-sided Wilcoxon rank-sum tests (*^∗^P <* 0.05*,^∗∗^ P <* 10*^−^*^2^*,^∗∗∗^ P <* 10*^−^*^3^). **c**) Distribution of DG counts (per vOTU) across environments. **d**) Proportion of DG across phage families within each environment. **e**) Distribution of DG across phage lifestyles and environments, expressed as percentages. **f**) Comparison of DG density (per kb) between regular phages (genome size *<* 175 kb) and jumbo phages (genome size *>* 200 kb) across biomes.

**Suppl. Fig. 3.**
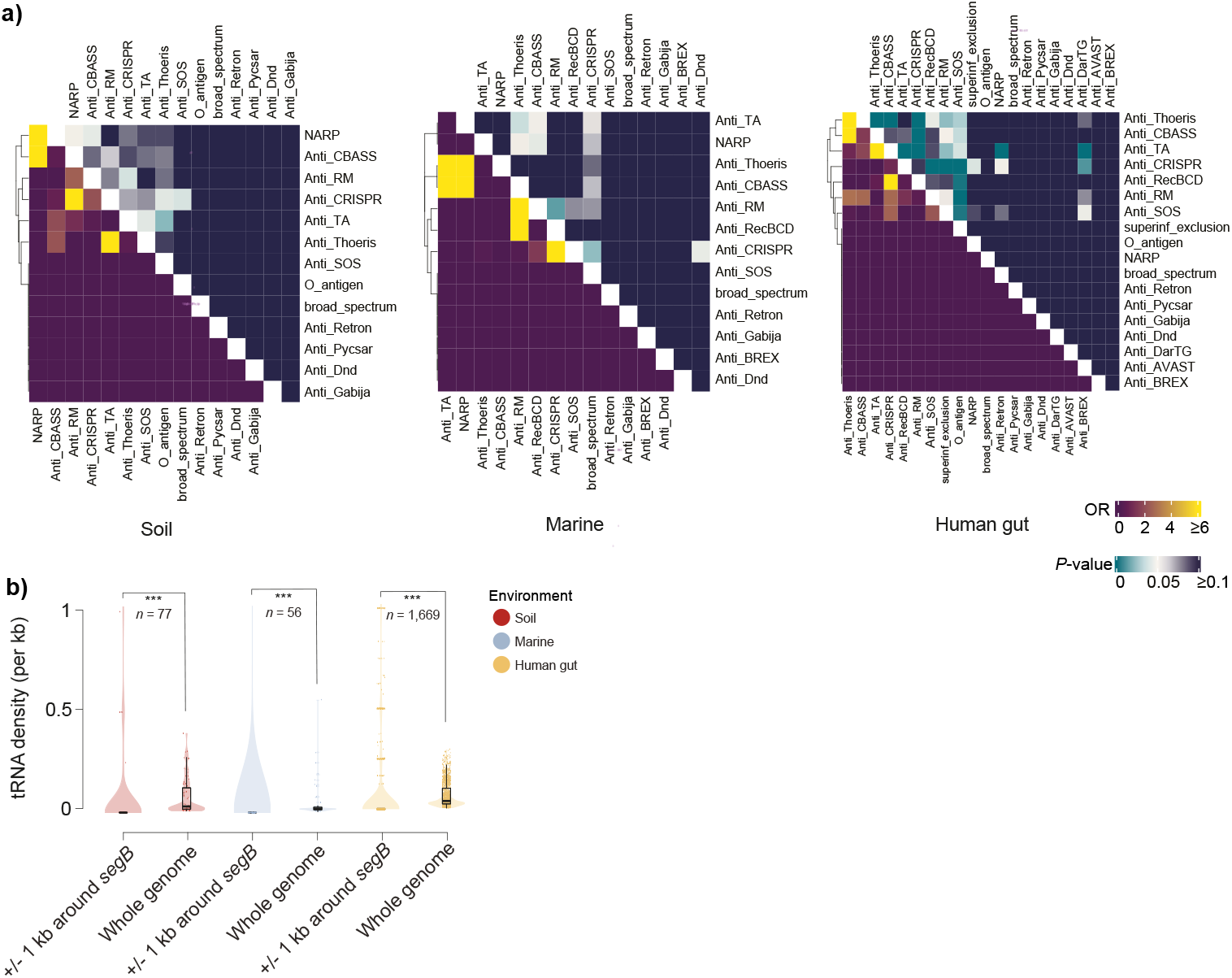
Colocalization across soil, marine, and human gut vOTUs. **a**) Odds ratio (OR) of co-localization (bottom heatmaps) and associated two-sided Fisher’s exact test *P* values (upper heatmaps) among counter-defense gene families, restricted to families with a co-localization frequency *≥*1% across soil, marine, and human gut vOTUs. **b**) Distribution of tRNA density (per kb) within +/-1 kb upstream and downstream of *segB* genes and across the whole viral genomes. Each dot represents a phage genome. Boxplots depict the interquartile range (25th - 75th percentiles), with the median indicated by a central black line. Whiskers extend to 1.5x the interquartile range, and outliers beyond tRNA densities = 1 were omitted for improved visualization. Statistical comparisons were performed using two-sided Wilcoxon rank-sum tests (*^∗∗∗^P <* 10*^−^*^3^). The number of vOTUs included in each category is indicated.

**Suppl. Fig. 4.**
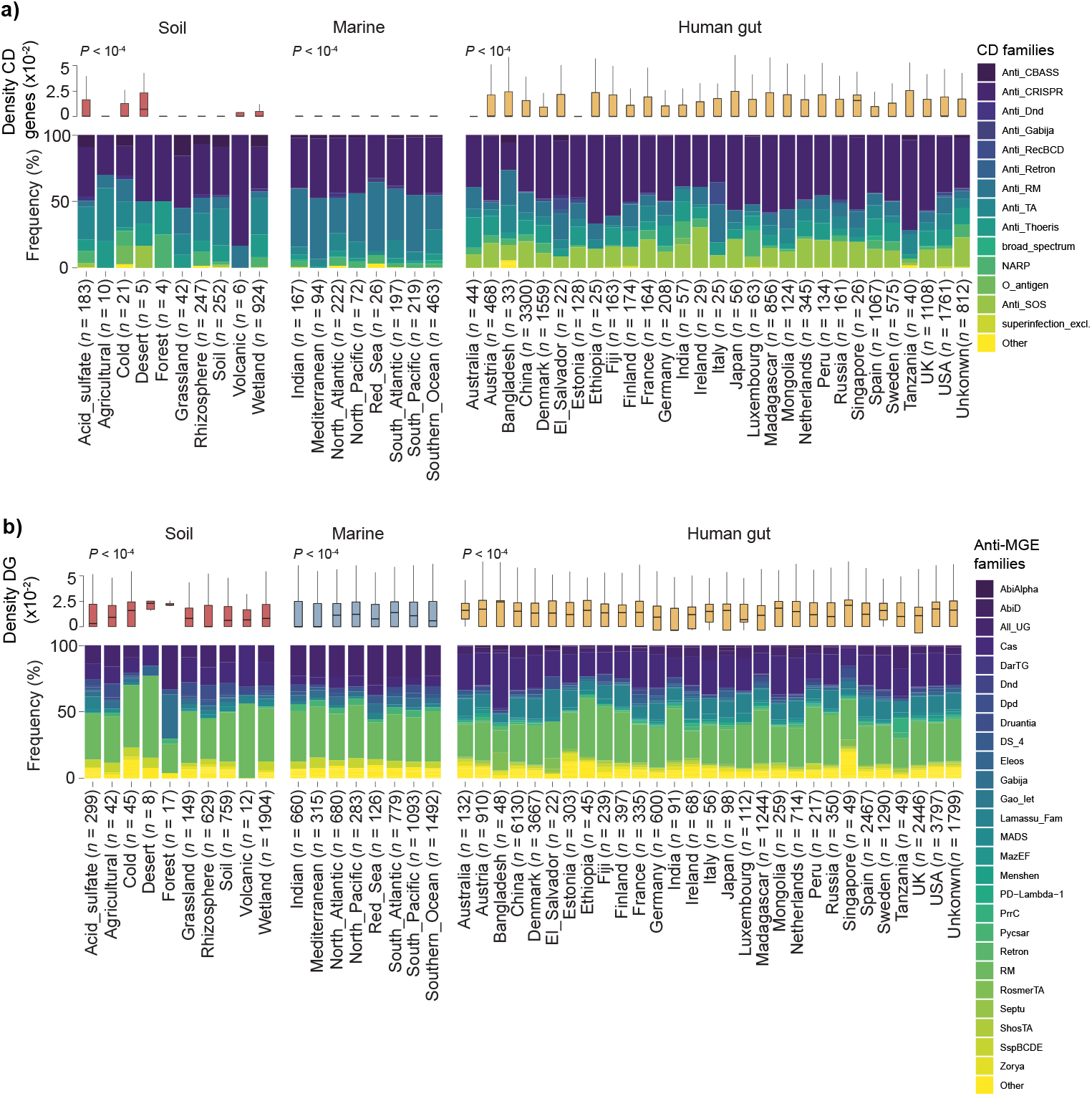
Interplay between phage biogeography, counter-defense (CD) gene abundance, and anti-MGE gene abundance. **a**) Density and frequency of CD genes across distinct ecological (soil, marine) and geographical (human gut) contexts. **b**) Density and frequency of anti-MGE genes across the same biomes. Boxplots depict the interquartile range (25th - 75th percentiles), with the median indicated by a central black line. Whiskers extend to 1.5x the interquartile range, and outliers were omitted for improved visualization. Statistical significance was assessed using a Kruskal-Wallis test. The number of vOTUs included in each category is provided in parentheses.

**Suppl. Fig. 5.**
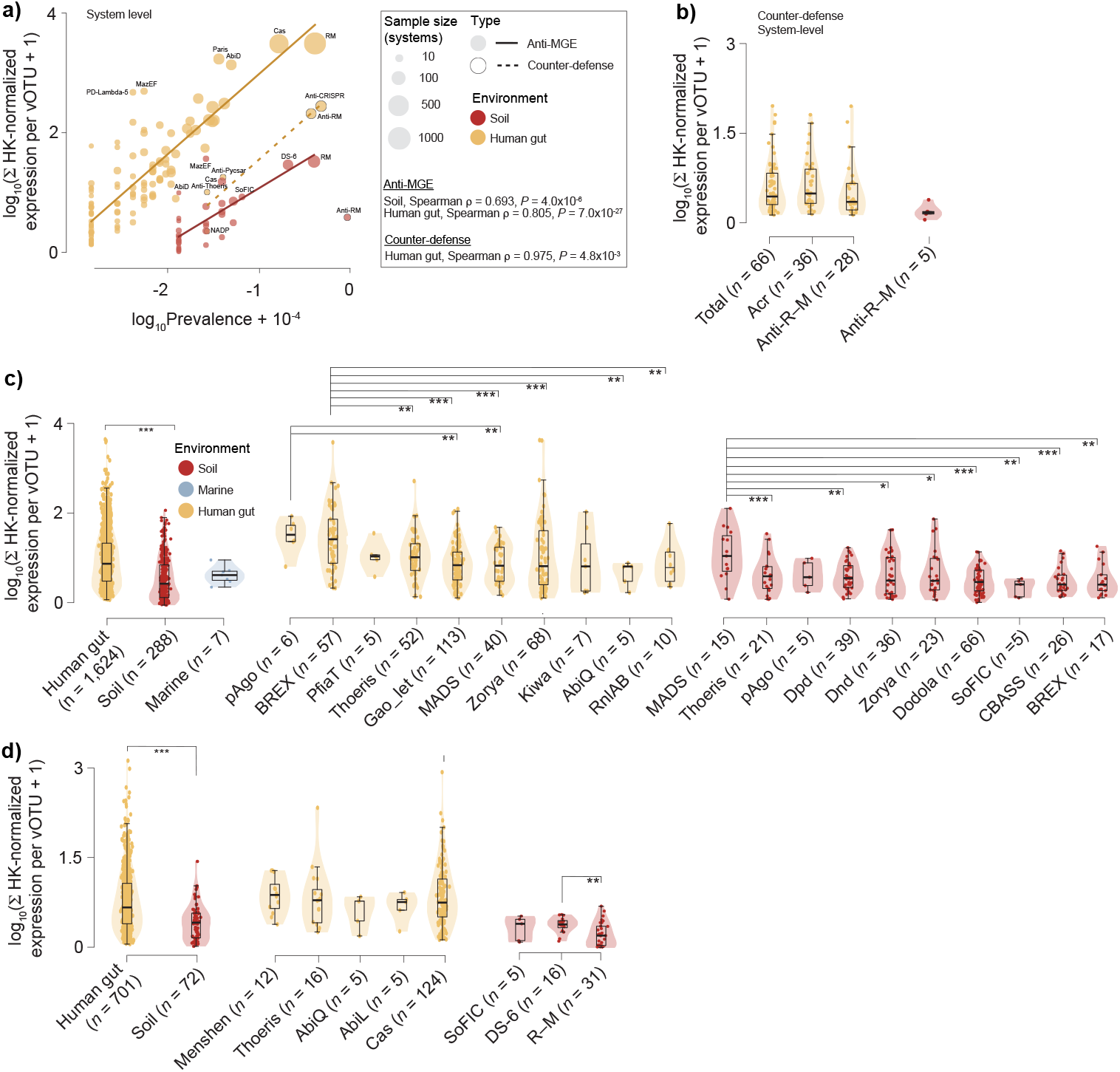
Interplay between counter-defense repertoires and their transcriptome. **a**) Relationship between immune system prevalence and sum of housekeeping (HK)-normalized expression per vOTU across complete anti-MGE (solid trend lines) and counter-defense (dashed trend lines) system families in soil (red) and human gut (yellow) samples. Each point represents one immune family sized by the number of contributing systems. Spearman correlation coefficients and *P* values are shown for each immune class and environment. Marine was devoid of complete systems passing filters. **b**) Summed HK-normalized expression per vOTU of counter-defense systems carried by soil (red), and human gut (yellow) vOTUs. *n* corresponds to the number of vOTUs per category, and only those with *n ≥* 5 are shown. **c**) Summed HK-normalized expression per vOTU of anti-MGE genes carried by soil (red), marine (blue), and human gut (yellow) vOTUs (left), or split by the top 10 most-expressed families for soil and human gut vOTUs. Not enough statistical power was obtained to represent marine samples when split by family. **d**) Summed HK-normalized expression per vOTU of anti-MGE systems carried by soil (red), and human gut (yellow) vOTUs (left), or split by the most-expressed families for soil and human gut vOTUs. *n* corresponds to the number of vOTUs per category, and only those with *n ≥* 5 are shown. Statistical comparisons were performed using two-sided Wilcoxon rank-sum tests (*^∗^P <* 0.05*,^∗∗^ P <* 10*^−^*^2^*,^∗∗∗^ P <* 10*^−^*^3^).

**Suppl. Fig. 6.**
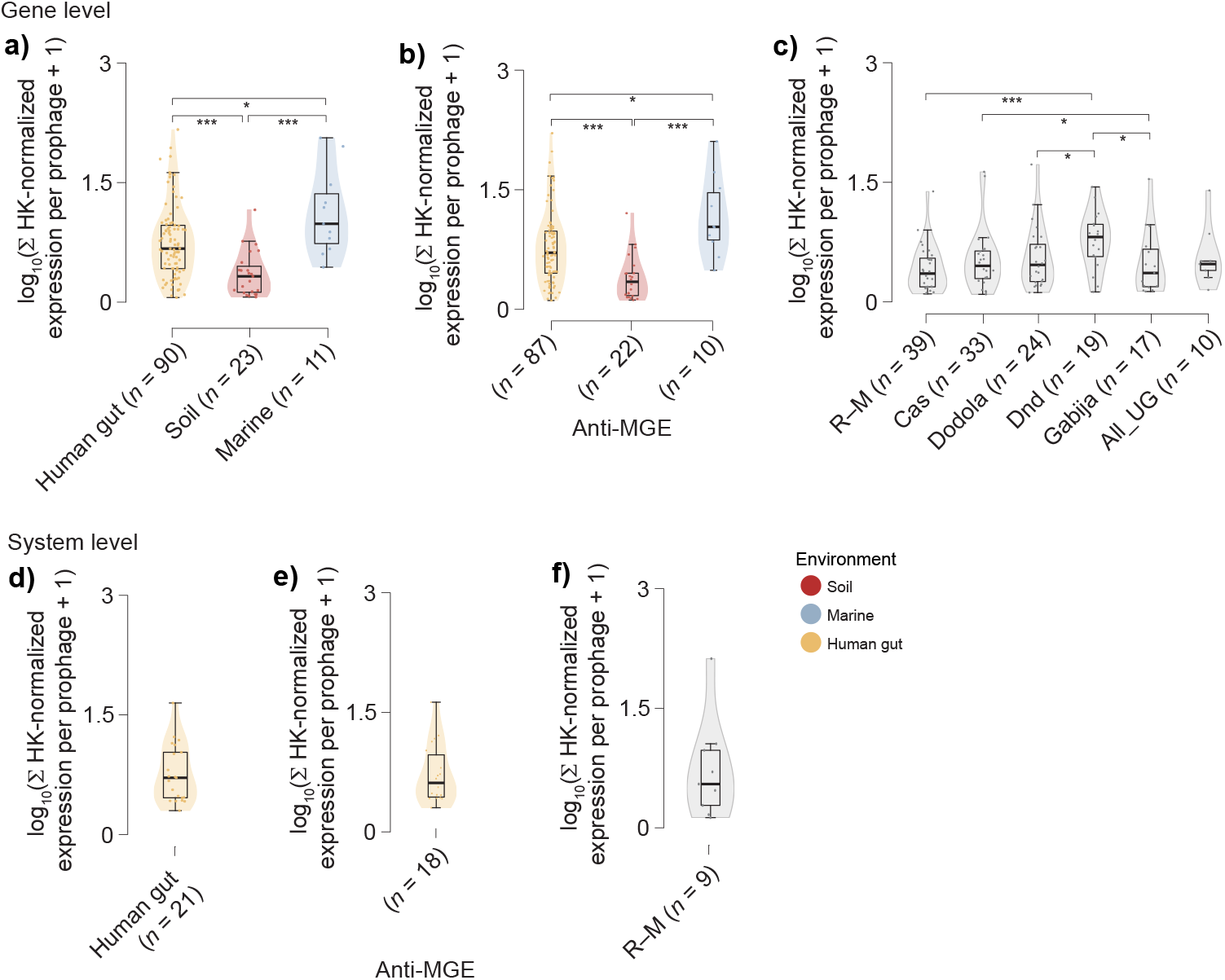
Transcriptomic profiling of prophage-associated immune repertoires in bacterial MAGs reconstructed from the same environments as the vOTUs. **a**) Summed housekeeping (HK)-normalized expression per prophage of counter-defense + anti-MGE defense genes carried by soil (red), marine (blue), and human gut (yellow) bacterial MAGs. *n* corresponds to the number of prophages per category, and only those with *n ≥* 5 are shown. **b**) Same as **a**) but considering only anti-MGEs. **c)** Summed HK-normalized expression of anti-MGE genes (split by anti-MGE family) carried by bacterial MAG prophages. **d**) Summed HK-normalized expression per prophage of counter-defense + anti-MGE defense systems carried by human gut bacterial MAGs. **e**) Same as **d**) but considering only anti-MGE systems. **f**) Summed HK-normalized expression of R–M systems carried by bacterial MAG prophages. Statistical comparisons were performed using two-sided Wilcoxon rank-sum tests (*^∗^P <* 0.05*,^∗∗^ P <* 10*^−^*^2^*,^∗∗∗^ P <* 10*^−^*^3^).

**Suppl. Fig. 7.**
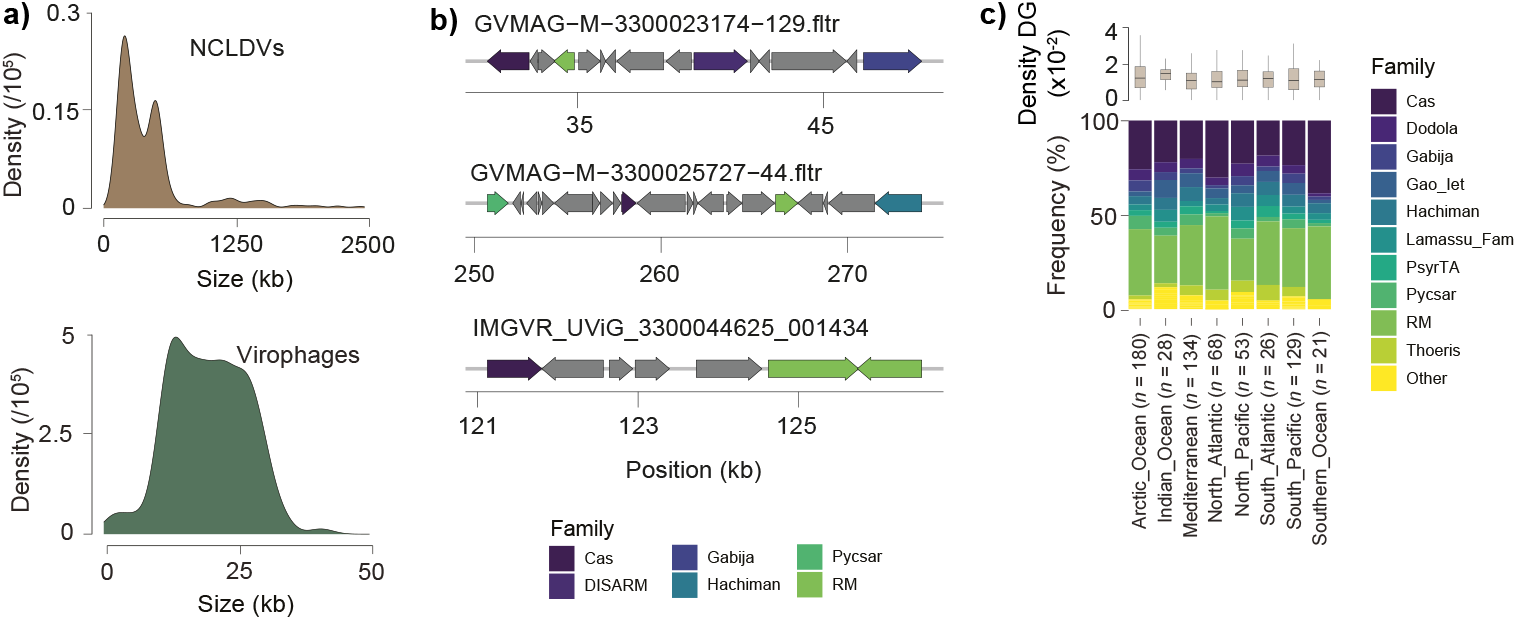
Clustering and biogeographical distribution of immune modules in NCLDVs. **a**) Size distribution patterns of the virophage and NCLDV dataset. **b**) Representative examples of immune module clustering in NCLDVs. **c**) Density and frequency of anti-MGE and counter-defense gene families in NCLDVs recovered from distinct marine regions. Boxplots depict the interquartile range (25th - 75th percentiles), with the median indicated by a central black line. Whiskers extend to 1.5x the interquartile range, and outliers were omitted for improved visualization. Statistical significance was assessed using a Kruskal-Wallis test. The number of NCLDV genomes included in each category is provided in parentheses.

## Notes

### Competing Interest Statement

The authors have declared no competing interest.

### Summary of Updates

We have now added additional analyses to the manuscript, including: i) the expression landscape of immune genes carried by prophages identified in bacterial MAGs from the same biomes from which the vOTUs were recovered (new Suppl. Figs. 6 a-f); ii) a comparison of counter-defense gene densities between conventional phages and jumbo phages (new Suppl. Fig. 2f); and iii) an analysis of the distinct patterns of segB-tRNA colocalization observed in our large-scale dataset (new Suppl. Fig. 3b).

